# RNAcmap: A Fully Automatic Method for Predicting Contact Maps of RNAs by Evolutionary Coupling Analysis

**DOI:** 10.1101/2020.08.08.242636

**Authors:** Tongchuan Zhang, Jaswinder Singh, Thomas Litfin, Jian Zhan, Kuldip Paliwal, Yaoqi Zhou

## Abstract

**Motivation:** The accuracy of RNA secondary and tertiary structure prediction can be significantly improved by using structural restraints derived from evolutionary or direct coupling analysis. Currently, these coupling analyses relied on manually curated multiple sequence alignments collected in the Rfam database, which contains 3016 families. By comparison, millions of non-coding RNA sequences are known. Here, we established RNAcmap, a fully automatic method that enables evolutionary coupling analysis for any RNA sequences. The homology search was based on the covariance model built by Infernal according to two secondary structure predictors: a folding-based algorithm RNAfold and the latest deep-learning method SPOT-RNA.

**Results:** We show that the performance of RNAcmap is less dependent on the specific evolutionary coupling tool but is more dependent on the accuracy of secondary structure predictor with the best performance given by RNAcmap (SPOT-RNA). The performance of RNAcmap (SPOT-RNA) is comparable to that based on Rfam-supplied alignment and consistent for those sequences that are not in Rfam collections. Further improvement can be made with a simple meta predictor RNAcmap (SPOT-RNA/RNAfold) depending on which secondary structure predictor can find more homologous sequences. Reliable base-pairing information generated from RNAcmap, for RNAs with high effective homologous sequences, in particular, will be useful for aiding RNA structure prediction.

**Availability and implementation:** RNAcmap is available as a web server at https://sparks-lab.org/server/rnacmap/) and as a standalone application along with the datasets at https://github.com/sparks-lab-org/RNAcmap.

## INTRODUCTION

RNA structures are the foundations for their diverse functional roles ranging from catalysis, cell-signalling, to transcriptional regulation (1). Determining RNA structures by traditional experimental techniques such as X-ray crystallography, nuclear magnetic resonance, and cryogenic electron microscopy is costly and time-consuming. In fact, only 3% or 99 of 3016 RNA families from Rfam (2) have experimentally solved structures and the number of solved RNA-only structures per year stay the same for the past two decades (50∼80/year). By comparison, the number of noncoding RNAs collected in RNAcentral has doubled from 8 million in 2015 to 16 million in 2019(3, 4). The fast-increasing gap between the number of noncoding RNA sequences and the number of experimentally-solved structures makes computational approaches highly desirable.

Computational RNA structure predictions were evaluated by RNA puzzles(5–7), which were blind experiments in RNA 3-D structure prediction, similar to Critical Assessment of Structure Prediction (CASP) for blind protein structure prediction (8–10). Results of recent three rounds of RNA puzzles showed that predicting near-native models (RMSD <10 Å) remained challenging for most current methods (5–7). However, there is a significant improvement in ab initio protein structure prediction in CASP13 (11), largely due to employing significantly improved prediction in protein contact maps as the restraints for 3D structure prediction. Prediction of protein contact map was improved by increasingly accurate mutational coupling analysis due to the fast-expanding protein sequence database and more powerful, deep contextual learning for enhancing coupling signals (8, 12).

Contact maps inferred from mutational coupling have been demonstrated to improve RNA secondary (13, 14) and tertiary structure prediction (15–17). However, RNAalifold (13) relies on mutual information that is unable to separate direct and indirect coupling (18), while more accurate evolutionary coupling (16) or direct coupling (17) analysis relied on homologs identified in the Rfam database, which has 3016 families only as of January 2019. This limitation prevents a wide application of evolutionary coupling for RNA secondary and tertiary structure prediction.

An accurate mutational coupling analysis requires a large number of sequence homologs. RNA homology search is a challenging problem because most structural RNAs are known to preserve the secondary structure rather than the primary sequence(19). The covariance-model-based search enabled by Infernal was shown to outperform both sequence-based and profile HMM-based methods with very high sensitivity and specificity, as it incorporates information from both sequence and secondary structure(20). Recent studies on genome-wide search for pseudoknotted noncoding RNA(21, 22) and comparison of RNA multiple sequence alignment tools(23) confirmed the state-of-the-art performance of Infernal.

The purpose of this work is to develop a fully-automatic method (RNAcmap) for RNA evolutionary coupling analysis that is not limited to RNA homologous sequences identified in Rfam. RNAcmap first employs BLAST (24) to perform an initial homolog search from the NCBI nucleotide database. The resulting homologous sequences and the predicted secondary structure are then employed for building the covariance model for the second-round search by Infernal (25), the same tool employed in Rfam to facilitate the comparison. Unlike Rfam that utilizes experimentally validated secondary structures or consensus prediction, RNAcmap employs a folding-based algorithm RNAfold (26) or a recent deep-learning-based method SPOT-RNA (27) for secondary structure prediction to ensure that the method is fully automatic. The resulting multiple sequence alignment from the second-round search is then employed for evolutionary coupling analysis to yield base-pairing and distance-based contact maps. Three methods for evolutionary coupling analysis are examined (GREMLIN, plmc and mfDCA) (15, 17, 28).

We showed that the resulting contact maps from RNAcmap (SPOT-RNA) are comparably accurate to those based on Rfam-aligned homologous sequences for those sequences in Rfam. More importantly, RNAcmap can yield similarly accurate base-pairing results for those sequences that are not curated in Rfam. RNAcmap should be useful for improving RNA secondary and tertiary structure prediction.

## MATERIALS AND METHODS

### The RNAcmap Pipeline

The RNAcmap pipeline provides two rounds of the search for RNA homologs. The first round of search is based on a purely sequence-based search tool, BLAST-N (24), against the NCBI nucleotide database. The parameters of `-evalue 0.001, line-length 1000`, and `-task blastn` are used in this round. The homologous sequences are obtained by an e-value cut off of 0.001. Then a covariance model is built using cmbuild from Infernal (25) with the above-obtained homologous sequences and predicted secondary structure from a pre-chosen secondary-structure prediction tool as input. Afterwards, the covariance model is employed to perform the second round of search against the NCBI database by cmsearch of Infernal with parameters `--incE 10.0`. The new updated homologous sequences (based on a cut off of E-value 10.0) and their alignment are used for evolutionary coupling analysis with a chosen tool. We employed a cut off of 10.0 for E-value in cmsearch as Sun et al in order to include more homologs with low sequence identity(29).

### RNA Secondary Structure Prediction Tool

We employed either a folding-based algorithm RNAfold (26) or our recently developed deep-learning method SPOT-RNA (27) for secondary structure prediction. SPOT-RNA improves prediction of secondary structure over existing folding-based algorithms, not only in canonical but also in non-canonical and non-nested (pseudoknot) base pairs. The improvement is the largest for non-nested and non-canonical base pairs (27).

### Evolutionary Coupling Analysis Tool

Several tools are employed for final evolutional coupling analysis. GREMLIN(28) is obtained from https://github.com/sokrypton/GREMLIN_CPP. The method is based on the pseudo-likelihood inference of direct coupling analysis with L2 regularization. GREMLIN is run with the recommended parameter for RNA `(-alphabet rna -gap_cutoff 1.0 -lambda 0.01 -eff_cutoff 0.8 -max_iter 100)`. The plmc method was obtained from https://github.com/debbiemarkslab/plmc. It employed similar pseudo-likelihood to infer the parameters in the DCA model (17). We utilized the recommended parameters for RNA `(-a -.ACGU -le 20 -lh 0.01 -m 50)`. The mfDCA method was obtained from http://dca.rice.edu/portal/dca/download. It employed the inverse covariance matrix to infer the coupling parameters(15, 18). An additional Average Production Correction (APC) (30) was applied to the original mfDCA to be consistent with other DCA methods.

### Evolutionary coupling analysis using Rfam-supplied multiple sequence alignment

RNAcmap can provide evolutionary coupling analysis based on a fully automatic search for homologous sequences. The Rfam database consists of 3016 manually curated RNA families (RFAM version 14.1) (2, 25). Thus, evolutionary coupling analysis based on multiple sequence alignment provided in Rfam can be considered as an upper limit for comparing with RNAcmap. This is because RNA secondary structures may be experimentally validated or from consensus prediction in Rfam-supplied alignment.

### Datasets

We downloaded a total of 4528 structures containing 6294 RNA chains from the Protein Databank. Among them, 4281 RNA chains were selected with sequence length between 32 and 500. Using cmfind and Rfam database (version 14.1)(2, 25), these chains were further split into two sets: 3182 RNA chains were mapped to existing Rfam families and 1099 RNA chains were not mapped to any Rfam families. The majority of structure-mapped Rfam families (77%, 2461 of 3182) are `tRNA, 5S rRNA` and 5.8S `rRNA`. These two sets were further reduced by limiting to X-ray-determined structures with resolution better than 3.5 Å and clustered by cd-hit-est(31, 32) with sequence identity cut off of 0.8.

For those RNAs mapped to known Rfam families, we required the minimal aligned length to be 80% of the RNA length and the aligned length of the Rfam covariance model to be greater than 50%. If one RNA was aligned to two or more Rfam families, the family with the highest score (or the lowest E-value) was taken as the correctly aligned family. Finally, for each family, we selected one RNA. The final Rfam set contained 43 structured, non-redundant RNAs.

For those RNAs that were not mapped to Rfam families, we required the minimal number of base pairs to be 10, which is the lowest number of base pairs in the Rfam set. This non-Rfam set contained 117 non-redundant RNAs.

We used X3DNA-DSSR(33) to annotate the secondary structure from the PDB structure and bpRNA (34) to annotate various base pair types and stem regions for both predicted and reference secondary structure. Briefly, a stem is defined as a helix consisting of only canonical Watson-Crick and wobble pairs with a continuous backbone. An isolated canonical basepair is defined as a basepair without stacking interaction or not belonging to a stem. Pseudoknots exist when two non-nested base pairs (i, j) and (a, b) satisfy (i<a<j<b). Pseudoknot base pairs are those pairs that require the least to remove to yield pseudoknot-free secondary structure.

### Performance Measures for Base Pair Prediction

The performance of a single RNA was evaluated by the sensitivity (SN=TP/(TP+FN)), precision (PR=TP/(TP+FP)) and F1-score (the harmonic mean of sensitivity and precision, F1=2SN*PR/(SN+PR)) using top L/6, L/4, L/2 and L predicted pairs. Here, TP, TN, FP and FN are true positives, true negatives, false positives and false negatives, respectively. Only non-local pairs were evaluated (|i-j|>4, i and j are the sequence positional indices of the nucleotides).

### Performance Measures for Stem Prediction

Because evolutionary or direct coupling only captures a few base pairs within a stem with the strongest co-mutation signals, it is necessary to evaluate the performance on the stem level, rather than on the base-pair level. This is because it is more important to capture all the stems for the overall correct topology, rather than obtain most base pairs in some stems but not even a single base pair in other stems. To evaluate the performance on the stem level, we defined that a stem is correctly predicted if one or more base pairs within the stem are correctly predicted.

### Performance Measures for Tertiary-contact Prediction

Here, we defined non-hydrogen-bonded tertiary contacts if a) the nearest-heavy atom distance between two nucleotides is less than 8Å and b) these two nucleotides are not adjacent to the existing base pairs. This definition follows the work by De Leonard et al(15). Tertiary contacts were evaluated after removing base pairs and two nucleotides neighbouring to the base pairs. The overall performance was also evaluated by F1-score, sensitivity, and precision.

## RESULTS

### Comparison Between Evolutionary Coupling Tools for RNAcmap (RNAfold)

RNAcmap requires a secondary structure prediction tool for building the covariance model and an evolutionary coupling tool for predicting contacting pairs. In RNAcmap (RNAfold), we employed RNAfold for secondary structure prediction and examine how different evolutionary coupling tools would impact the outcome of contact prediction. Figure 1 compares F1-score, precision, and sensitivity for base-pair prediction in the Rfam dataset by GREMLIN, mfDCA_apc and plmc, respectively. Results for top L/6, L/4, top L/2 and top L predictions are presented for 43 RNAs in the Rfam set. As expected, increasing the number of predictions from top L/6 to L leads to an increase in sensitivity but a decrease in precision. Top L/4 predictions reached the highest F1-score for all evolutionary coupling methods, suggesting that L/4 predictions have the optimal balance of sensitivity and precision.

**Figure 1.**
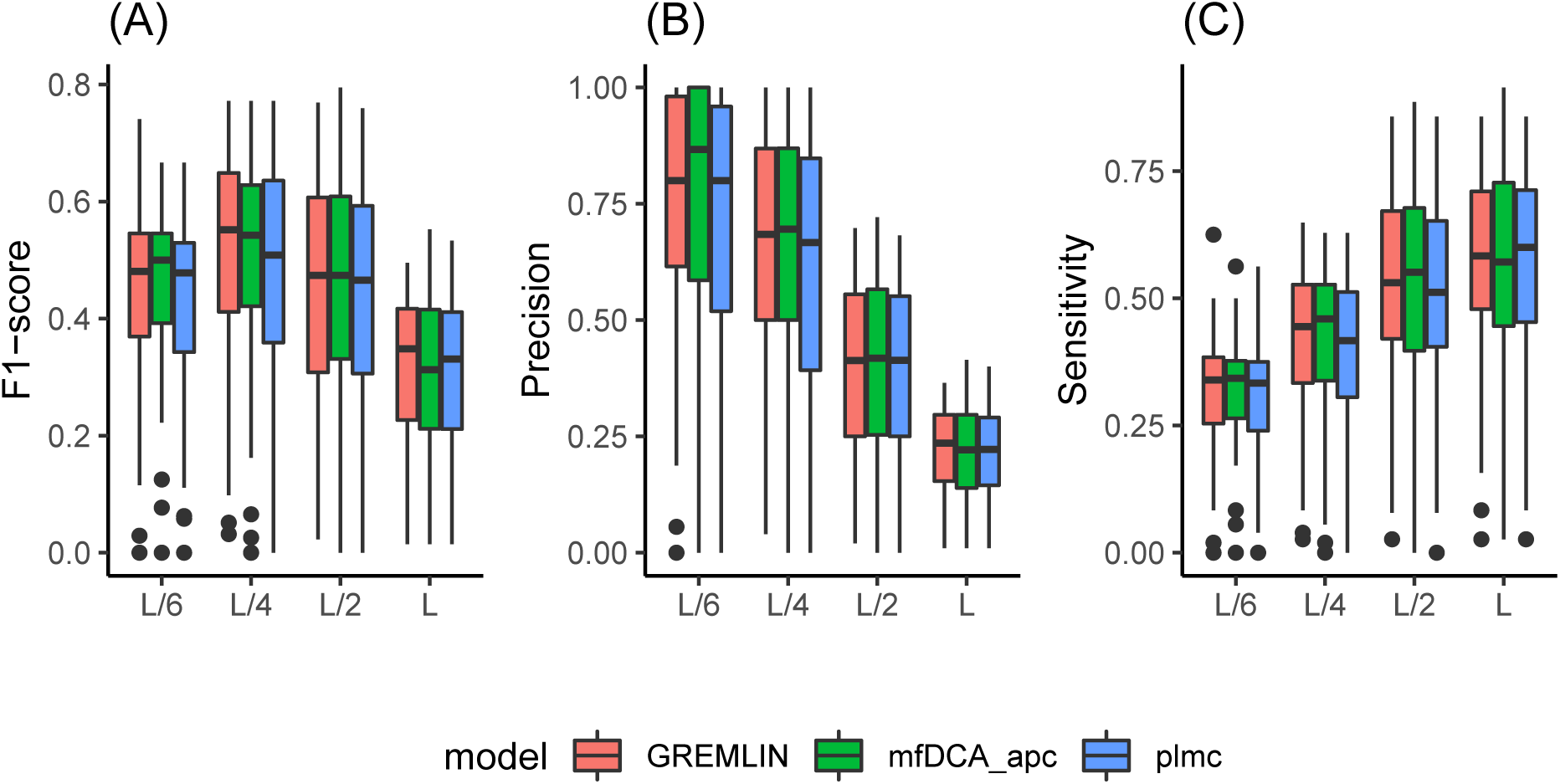
Boxplot of F1-score (A), Precision (B), and Sensitivity (C) of predicted base pairs by RNAcmap (RNAfold) based on three evolutionary coupling methods GREMLIN, mfDCA_apc and plmc, respectively, for 43 RNAs in the Rfam set.The distribution is shown in terms of median, 25th and 75th percentile with outlier shown by dots.

GREMLIN and mfDCA had comparable performance. For example, using the F1 score for top L/4 prediction as a measure, there was no statistically significant difference between GREMLIN and mfDCA (p-value=0.81, the paired t-test). Both, however, were statistically significant but only slightly better than plmc (p-value=0.005 for mfDCA better than plmc, p-value = 0.0007 for GREMLIN better than plmc). Because the difference between the three methods was small, GREMLIN will be used as the default for all subsequent analysis.

### Evolutionary Coupling Analysis from Rfam-supplied alignment and from the alignment generated from RNAcmap (RNAfold)

Using the same Rfam dataset, Figure 2 compares F1-score, precision, and sensitivity of base-pair prediction from evolutionary coupling analysis by GREMLIN with multiple-sequence alignment generated by RNAcmap (RNAfold) and supplied by Rfam, respectively. For a reference, the result based on the first-round BLAST-N search is also shown. BLAST-N-based alignment provides very poor (nearly random) prediction. Manually curated alignments from Rfam improves over automatic homology search and alignment given by RNAcmap (RNAfold). The improvement is observed for both sensitivity and precision. Rfam-alignment improves over RNAcmap (RNAfold) statistically significant for top L/6 predictions (p-value=0.02, paired t-test on F1-score of 43 RNAs), top L/4 predictions (p-value=0.01), and top L/2 predictions (p-value=0.04) but not top L predictions (p-value=0.06).

**Figure 2.**
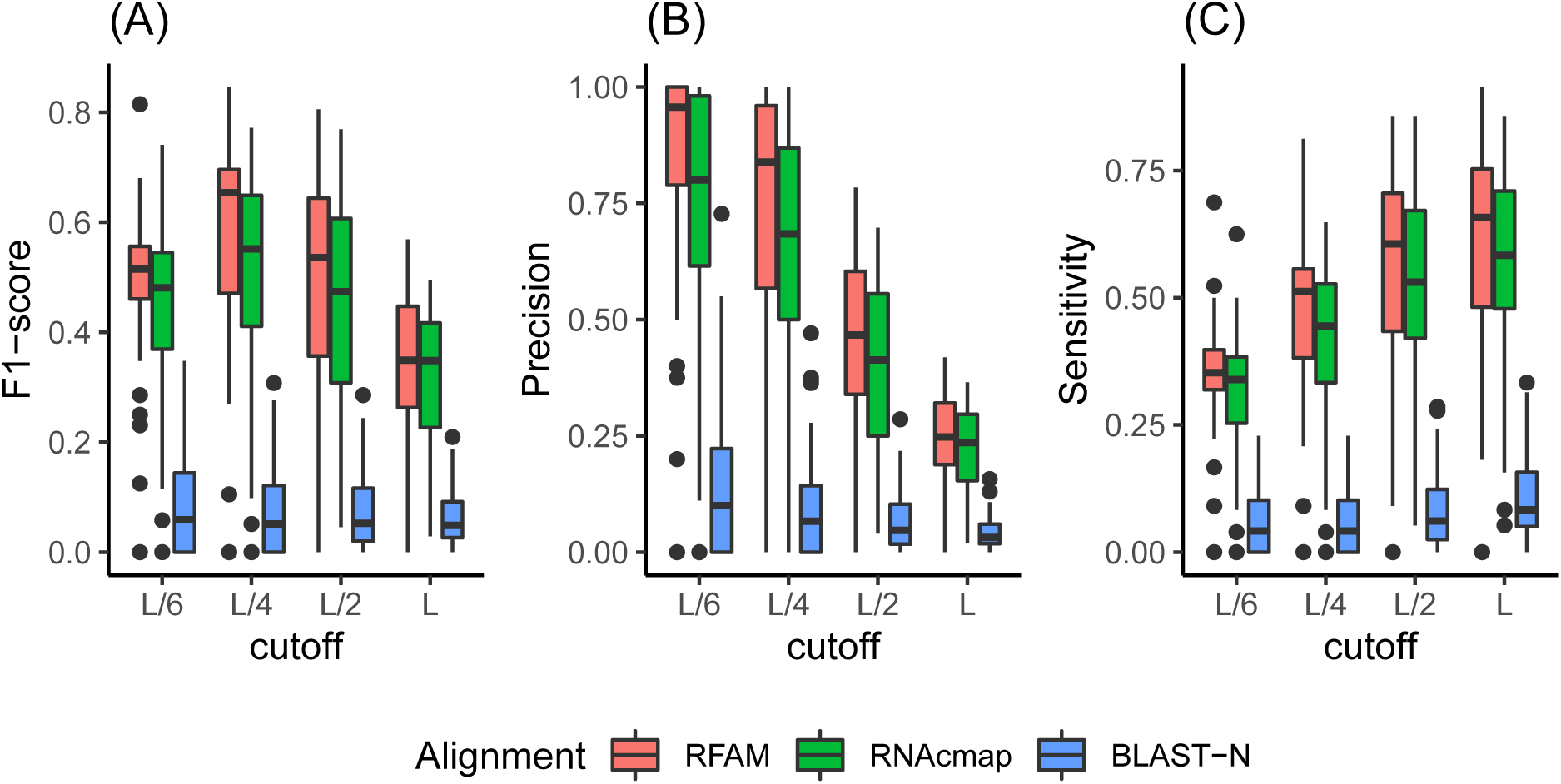
Boxplot of F1-score (A), Precision (B), and Sensitivity (C) by Rfam-supplied alignment in comparison to RNAcmap (RNAfold) and BLAST-N for 43 RNAs in the Rfam set. The distribution is shown in terms of median, 25th and 75th percentile with outlier shown by dots. All employed GREMLIN for base-pair prediction

Figure 3 further examine the ability to predict tertiary contacts by comparing the results of Rfam-based alignment, RNAcmap (RNAfold), and Blast-N. Performance for all methods is poor with <5% in median precision and <2% in median sensitivity for all top L/6, L/4, L/2 and L predictions although Rfam-based alignment and RNAcmap (RNAfold) are significantly better than Blast-N (with p-value<0.0002 for all cases). Thus, here and hereafter we will focus on base-pair prediction only.

**Figure 3.**
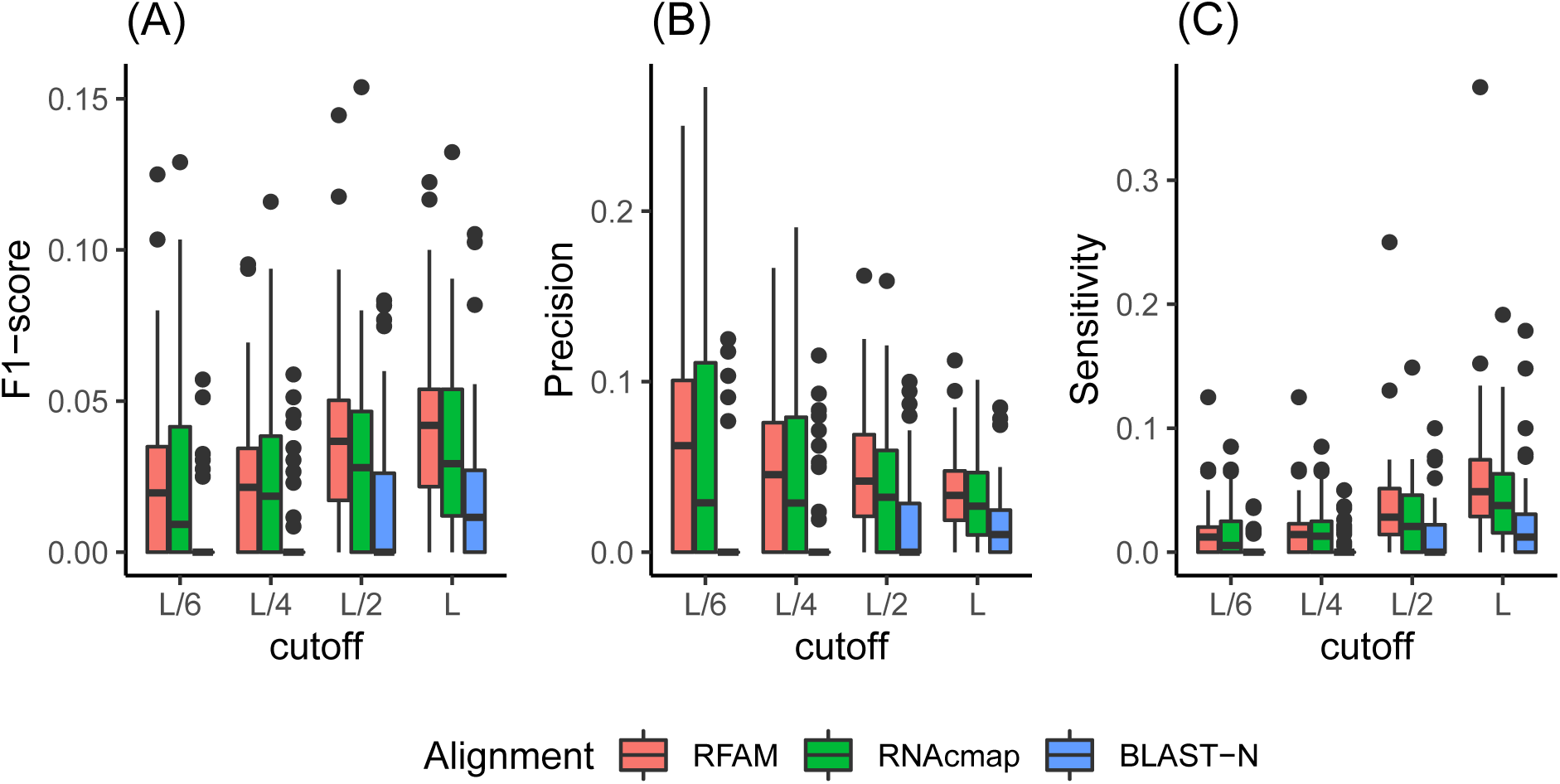
As in Figure 2 but for distance-based tertiary contact prediction

### Beyond Rfam sequences

Although Rfam-based alignment has a slight edge in performance for top L/4 predictions, in particular, one advantage of RNAcmap (RNAfold) is that it can predict sequences which are not in the Rfam collection. Figure 4 illustrates F1-score for the Top L/4 predictions as a function of the number of effective homologous sequences, normalized by sequence length (Neff/L). While the Rfam-based alignment improves over RNAcmap (RNAfold) for the Rfam set, the performance of RNAcmap (RNAfold) for 117 RNAs in the non-Rfam set is nearly the same as that for 43 RNAs in the Rfam set. All showed a trend of improved prediction with increased Neff/L. Particularly, reasonably accurate predictions (F1-score>0.5) are made for Neff/L>1 for 21 of 27 RNA sequences (78%) for the Rfam set using Rfam alignment, 15 of 20 RNA sequences (75%) for the Rfam set using RNAcmap (RNAfold) alignment, and 16 of 19 RNA sequences (84%) for the non-Rfam set using RNAcmap (RNAfold) alignment. Two outliers with high Neff/L and low F1-score in Figure 4 were both resulted from poorly predicted secondary structures (red:6ASO:I, blue: 4QJD:B), which led to incorrect homologous sequences.

**Figure 4.**
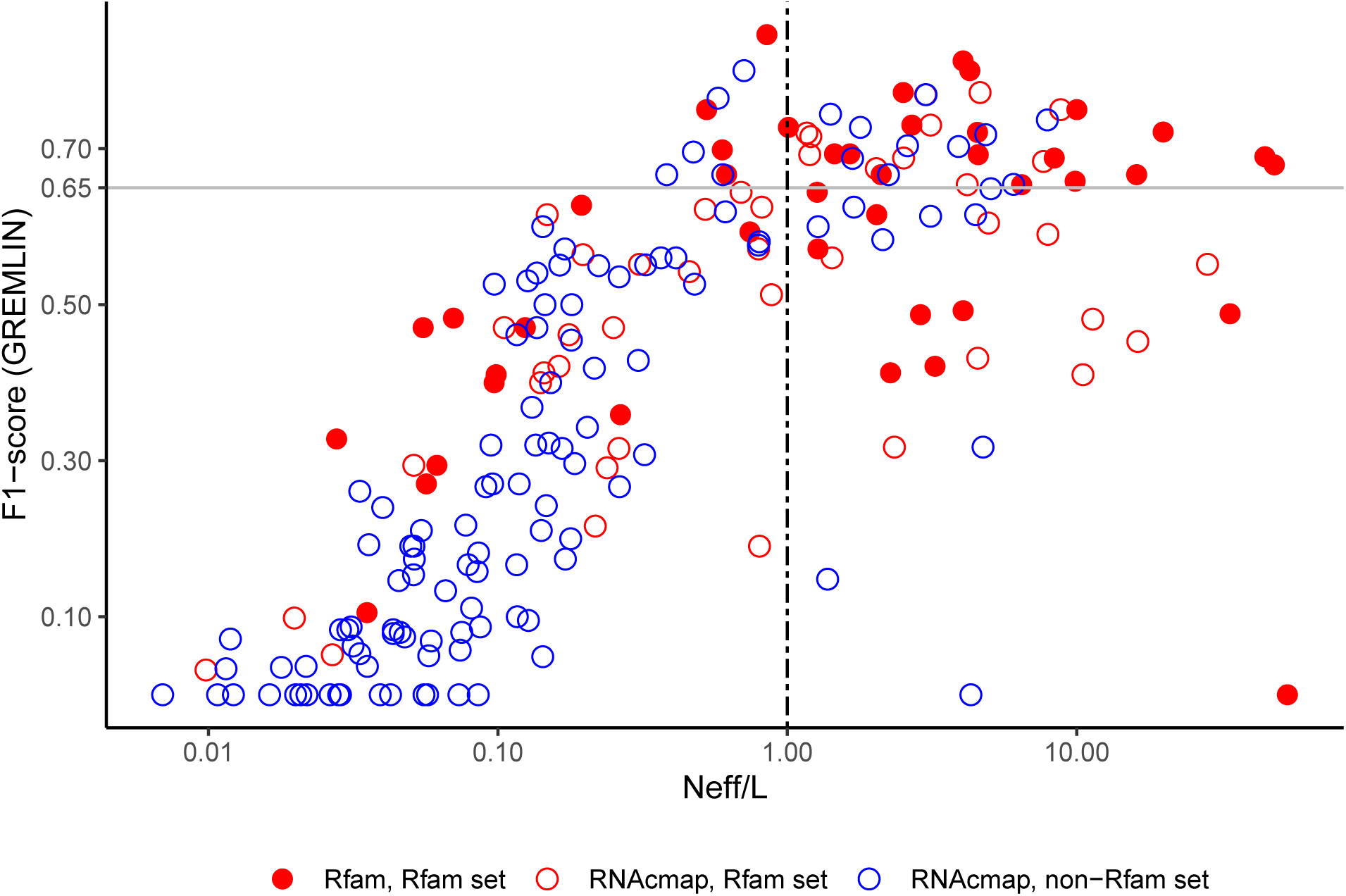
F1-score of base-pair prediction as a function of alignment effective size normalized by length (Neff/L). GREMLIN was used for evolutionary coupling analysis using top L/4 pair predictions by Rfam-supplied alignment (filled red circle) and RNAcmap (RNAfold, red circle) for the Rfam set, and RNAcmap (RNAfold, blue) for the non-Rfam set.

### The Effect of Secondary Structure Prediction on the Performance of RNAcmap

We compared the effect of using RNAfold and SPOT-RNA in the RNAcmap pipeline. To make a fair comparison, we excluded those RNAs in Rfam and non-Rfam sets with sequence similarity greater than 80% to any of RNAs in the SPOT-RNA training set. This sequence-identity cut off was the lowest cutoff allowed by cd-hit-est (31, 32)and employed previously for separating training and independent test sets (27, 35, 36). This leads to a total of 77 RNAs as a combined test set (21 in the Rfam set and 56 in the non-Rfam set).

Figure 5 compares the performance in F1 scores for the top L/4 predictions by RNAcmap (SPOT-RNA) and by RNAcmap (RNAfold), respectively. It is clear that RNAcmap (SPOT-RNA) offers a substantial improvement over RNAcmap (RNAfold). The former has 31 RNAs with higher F1-scores, compared to 18 RNAs with higher F1-scores by the latter. The median F1 score is 0.35 for RNAcmap (SPOT-RNA) and 0.29 for RNAcmap (RNAfold). The difference is statistically significant with p-value =0.037 for 18 RNAs with Neff/L >1 (paired t-test).

**Figure 5.**
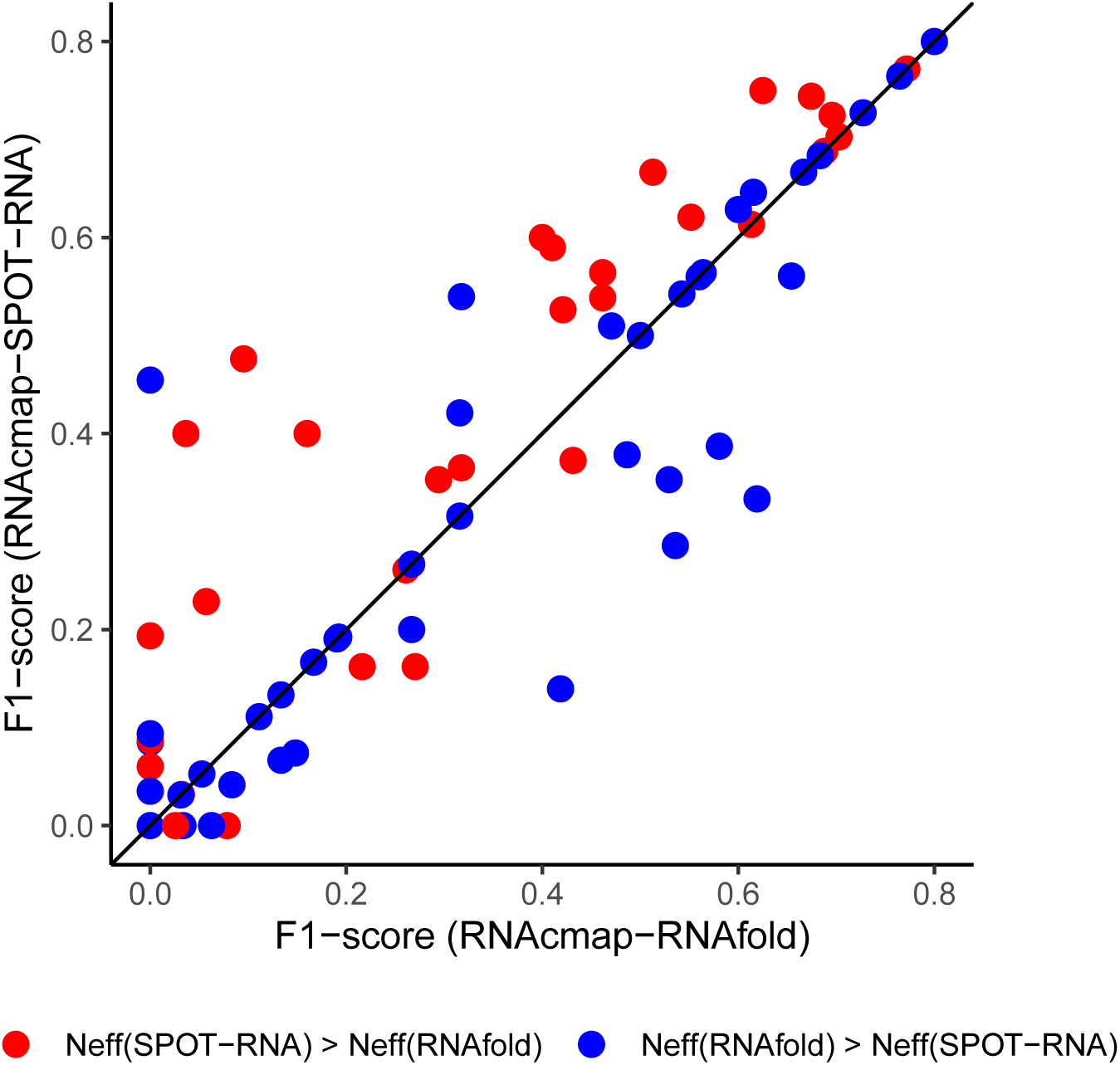
F1-score by RNAcmap with SPOT-RNA versus RNAcmap with RNAfold for 77 RNAs in a combined test set. RNAs with higher Neff by RNAcmap (SPOT-RNA) than RNAcmap(RNAfold) are shown in red (in blue, otherwise)

To further understand the source of the performance difference, Figure 5 also shows RNAs in different colours depending on whether Neff obtained from RNAcmap (SPOT-RNA) is higher or lower than that from RNAcmap (RNAfold). For many cases, the performance of RNAcmap is better for higher Neff/L, regardless if SPOT-RNA or RNAfold is employed. This suggests that it is possible to make a simple meta predictor RNAcmap (SPOT-RNA/RNAfold) depending on whether RNAcmap (SPOT-RNA) or RNAcmap (RNAfold) yields a higher Neff/L. This meta server would produce a median F1-score of 0.37 for the combined test set, compared to 0.35 by RNAcmap (SPOT-RNA) and 0.29 by RNAcmap (RNAfold). The difference is slightly better but not statistically significant with p-value <0.1 from RNAcmap (SPOT-RNA) but is significantly better with p-value <0.004 from RNAcmap (RNAfold). This meta server can achieve 14 of 18 RNA sequences (78%) with accurate predictions (F1-score>0.5) in the combined test set using RNAcmap alignment, compared to 10 by RNAcmap (RNAfold) or 13 by RNAcmap (SPOT-RNA). It should be noted that for 21 Rfam sequences in this combined set, the performance difference between contact maps from RNAcmap (SPOT-RNA) and those from Rfam-supplied alignments are statistically insignificant (p-value=0.28). The difference between contact maps from Rfam-supplied alignments and those from RNAcmap (RNAfold) is still significant (p-value=0.018) as in the full Rfam set.

One interesting question is how the base pairs obtained from evolutionary coupling differ from those directly predicted by secondary structure predictors. Figure 6A shows the results for 18 RNAs in the combined test set with Neff/L>1 as we considered that the evolutionary information is not reliable for Neff/L<1 as shown in Figure 4. We evaluated the performance on the base-pair level and on the stem-level. On the base-pair level, we examined different types of base pairs including canonical base pairs in helical regions, in nonhelical regions (unstacked, isolated single base pairs), noncanonical base pairs, and nested base pairs. RNAfold (or SPOT-RNA) can correctly predict more canonical base pairs in a helical region but less in nonhelical regions than evolutionary coupling by using the corresponding predicted secondary structure for search and alignment (i.e. RNAcmap (RNAfold) or RNAcmap (SPOT-RNA)). On the other hand, RNAcmap (RNAfold) significantly improves over RNAfold in predicting non-canonical base pairs and base pairs in pseudoknots, whereas SPOT-RNA is slightly better in predicting non-canonical base pairs and much better in predicting base pairs in pseudoknots than RNAcmap (SPOT-RNA). This is because RNAfold was not built for predicting pseudoknots or noncanonical base pairs whereas SPOT-RNA, a deep learning technique, was trained for predicting any base pairs including pseudoknots and noncanonical base pairs. Moreover, pseudoknot pairs are not employed by Infernal for building covariance models. What is more revealing is the evaluation at the stem level (Figure 6 last columns): RNAcmap (SPOT-RNA) reaches a consistently higher precision and sensitivity than SPOT-RNA (or RNAfold).

**Figure 6.**
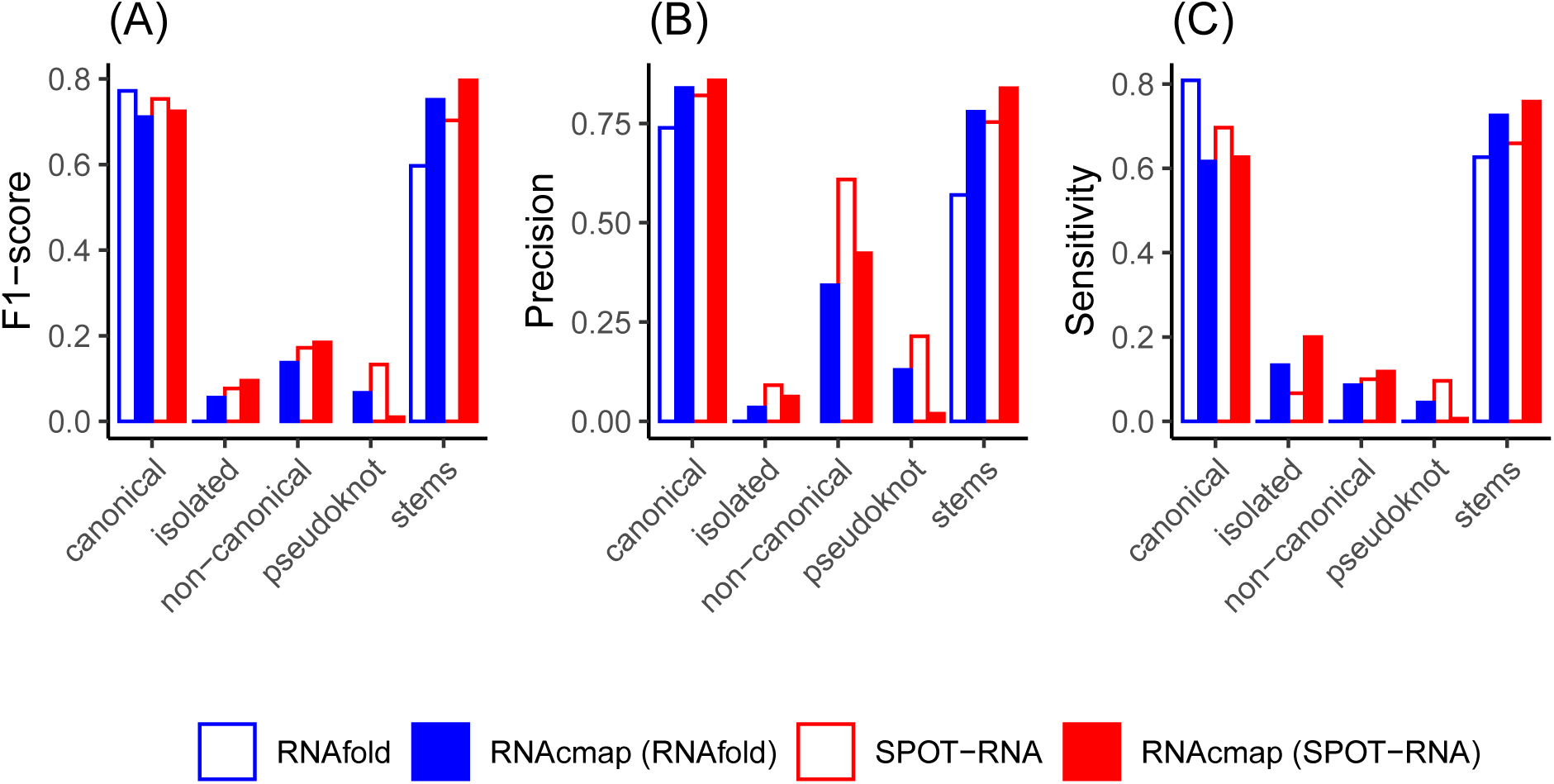
F1-score (A), Precision (B) and Sensitivity (C) of predicted base pairs reported according canonical, isolated canonical, noncanonical, pseudoknot base pairs and stems. The metrics are evaluated on 18 RNAs with “deep” RNAcmap alignment (Neff/L>1).

## Discussion

In this paper, we have established a fully automatic server RNAcmap that can predict contact maps directly from any given RNA sequences by homology search and evolutionary coupling analysis. The performance of RNAcmap (SPOT-RNA) or RNAcmap (SPOT-RNA/RNAfold) is comparable to that from manually curated Rfam sequences. More importantly, the performance is robust for those sequences not belonging to Rfam families. Thus, RNAcmap is expected to be useful to generate structural restraints for RNA secondary and tertiary structure prediction, as demonstrated previously(14–17).

It is found that the performance of RNAcmap is less dependent on the tools for evolutionary coupling analysis. The difference between GREMLIN, mfDCA_apc and plmc is small (Figure 1). This result is consistent with an independent study (23). However, the performance of RNAcmap is more strongly dependent on the secondary structure predictor (Figure 5). SPOT-RNA, that has more accurate secondary structure prediction, improves over RNA-fold in generating alignments that yielded improved contact map prediction. In particular, more stem regions were captured by using RNAcmap (SPOT-RNA) (Figure 6B), indicating more accurate topological connections in base pairing patterns. A simple meta predictor RNAcmap (SPOT-RNA/RNAfold) was established by using the secondary structure predictor that will yield a higher number of effective homologous sequences (Neff). This meta predictor further improves over RNAcmap (RNAfold). It is not entirely surprising as RNAfold (a folding-based algorithm) and SPOT-RNA (a deep-learning based method) are likely complementary to each other.

Contact map results for RNAs are different from those of proteins. For homologous sequence alignment, our results showed that sequence-only similarity search (BLAST-N) missed many homologous sequences with low sequence identity, resulting in poor prediction in the downstream evolutionary coupling analysis. Using a predicted secondary structure in the RNAcmap greatly expanded the coverage of homologous sequences, resulting in a much more accurate prediction. This is different in the case of protein homologous search, where sequence-only similarity is sufficient to capture most homologous sequences(37). Moreover, the contact maps for RNAs are dominated by the hydrogen-bonded base pairs. The accuracy for predicting distance-based tertiary contacts are only marginally better than random (Figure 3). This result is consistent with previous studies (15, 23).

However, not all base pairs are of equal importance when inferring the RNA structure. As showing in the comparison between RNAcmap and secondary structure predictors (SPOT-RNA(27), RNAfold(26)), RNAcmap-predicted base pairs are more enriched with isolated, pseudoknotted and non-canonical base pairs, which bring richer information for the overall topology of the RNA. Even for helical stem regions, RNAcmap predicts more stems than both SPOT-RNA and RNAfold, although the average number of predicted canonical base pairs within a stem is less than that of secondary-structure predictors. This is because evolutionary coupling can only capture the strongest signals that show marked difference in structural and functional stabilities between deleterious single and rescuing double mutations. In other words, the results from evolutionary coupling analysis offer a topology frame that can be further improved by a post processing method. Indeed, Zhe et al showed that a simple Monte-Carlo simulated annealing can recover nearly all base pairs of two ribozymes using pairing probabilities from mutational coupling analysis as a part of the energy function for folding secondary structure (14). A post processing model for RNAcmap is working in progress.

## Availability

RNAcmap is available as a webserver (https://sparks-lab.org/server/rnacmap/) and a standalone application (https://github.com/tcgriffith/RNAcmap)

## Acknowledgements

This work was supported by Australia Research Council DP180102060 to Y.Z. and K.P. We also gratefully acknowledge the use of the High-Performance Computing Cluster Gowonda to complete this research, and the aid of the research cloud resources provided by the Queensland CyberInfrastructure Foundation (QCIF). We gratefully acknowledge the support of NVIDIA Corporation with the donation of the Titan V GPU used for this research.

## Notes

### Competing Interest Statement

The authors have declared no competing interest.

